# From hyperinsulinemia to cancer progression: how diminishing glucose storage capacity fuels insulin resistance

**DOI:** 10.1101/2024.05.05.592630

**Authors:** Irina Kareva

## Abstract

Type 2 diabetes (T2D) is a complex metabolic disorder characterized by insulin resistance, hyperglycemia and hyperinsulinemia, with a quarter to half of people with T2D unaware of their diagnosis until the disease has reached advanced stages. T2D is associated with increased risk and worse prognosis of cardiovascular disease, cognitive decline, and cancer. Here we propose an updated framework for describing emergence of insulin resistance that precedes development of T2D. We show that diminishing capacity to store excess glucose can qualitatively capture the transition from normal to diabetic phenotype as captured by responses to oral glucose tolerance tests (OGTTs). We then show that an emerging tumor can either progress or regress depending on the metabolic environment of the host, consistent with experimental results of Hopkins et al. (2018), who showed that drug-induced transient diabetic phenotype, and specifically hyperinsulinemia, resulted in loss of therapeutic efficacy, and its reversal restored drug sensitivity and response to therapy. Given the prevalence of hyperinsulinemia in individuals with normoglycemia, addressing this condition emerges as a promising avenue to augment cancer therapy outcomes.

## Introduction

Type 2 diabetes (T2D) is a complex metabolic disorder that is characterized by insulin resistance and hyperinsulinemia, followed by progressive decline in the capacity of pancreatic beta cells to produce insulin ^1^. According to Ogurtsova et al. ^2^, in 2021 almost a quarter of adults in North America were unaware of having T2D, while in in West Africa, Western Pacific and Southeast Asia, this number can be over 50%. The incidence of pre-diabetes is estimated to be even higher ^3^, with nearly 70% of individuals with pre-diabetes expected to progress to overt diabetes within their lifetime ^4,5^.

In addition to being associated with higher incidence of complications, including nephropathy, chronic kidney disease and retinopathy, T2D is also associated with a higher incidence and worse prognosis of many other conditions, including cardiovascular disease, cognitive decline, and cancer ^6–9^. Intriguingly, it was recently demonstrated in animal models that a diabetic phenotype can cause resistance to certain types of cancer therapy, such as PI3K inhibitors. Specifically, in Hopkins et al. ^10^, the authors showed that administration of PI3K inhibitors to mice with implanted tumors caused emergence of both hyperglycemia and most importantly hyperinsulinemia, which in turn caused development of resistance to therapy. However, sensitivity to therapy was restored in all 12 tested tumor models when the authors corrected the diabetic phenotype through either administration of metformin, which reduces liver gluconeogenesis ^11^, through administration of sodium glucose cotransporter protein-2 (SGLT2) inhibitors, which increase glucose excretion in the urine, or via carbohydrate-restricted ketogenic diet. The authors confirmed that while elevated glucose may have contributed to emergence of resistance through providing cancer cells with extra substrate, it was correction of specifically hyperinsulinemia that caused restoration of sensitivity to therapy. In fact, it is well established that insulin can act as a powerful growth factor for cancer cells ^12–15^, and therefore normalizing baseline insulin levels diminished a growth signal, thereby allowing for the cytotoxic effects of the drug to outweigh the augmented growth of the tumor.

This result is particularly intriguing, since hyperinsulinemia can occur in individuals with normal blood glucose and is frequently undetected. In a classic study, Kraft et al. ^16^ analyzed the response of 3650 patients to an oral glucose tolerance test (OGTT), where a bolus of 100 g of glucose was administered to each person, and their blood glucose and insulin levels were assessed at baseline, 30, 60, 120 and 180 minutes ^16^; the analysis was subsequently re-done on a larger patient dataset by Crofts et al. ^17^ to address technical limitations of the original paper. Five different patterns of glucose-insulin dynamics were identified. Pattern I (norm), with fasting insulin levels maintained below 30 uIU/L, with a peak occurring at 30-60 minutes, followed by return to baseline by 180 minutes; the pattern of postprandial (after the meal) glucose increase and return to baseline is normal. Pattern II described individuals with still normal glucose dynamics but delayed return to baseline of insulin levels, suggesting that more insulin was becoming needed to stabilize glucose levels. Pattern III showed delayed insulin peak and higher postprandial glucose levels, as well as longer return to baseline. Finally, Kraft IV pattern revealed both higher postprandial levels of glucose, longer return to baseline and higher baseline levels of both insulin (over 50 uIU/L) and glucose, consistent with T2D; for these individuals, insulin production from the pancreas was not sufficient to lower blood glucose levels to normal levels. (Notably, Pattern V was also identified, where insulin levels never rose above 30 uIU/L, while the blood glucose levels were very high; this pattern is more reminiscent of type 1 diabetes (T1D) and is omitted in the subsequent analysis presented in this work, which is aimed at simulating emergence of insulin resistance, leading to T2D.) Critically, Crofts et al. ^17^ identified that over half of participants with normal glucose tolerance showed hyperinsulinemia despite normal glucose clearance ^17^, indicating that metabolic dysregulation occurs significantly prior to it being detectable on blood glucose tests.

The classical paradigm for understanding emergence of insulin resistance (IR) is through loss of sensitivity to signaling between the hormone insulin and its receptor. A classical metaphor used to describe this interaction is “lock and key”, where insulin (the “key”) binds to a lock (“insulin receptor”), causing translocation of an insulin-dependent glucose transporter, such as GLUT4, allowing glucose to enter the cell. If such an interaction is faulty (the metaphor of a “jammed lock”, in which the key cannot turn), then glucose cannot enter the cells and remains in the blood ^18^. An alternative mechanism to understand the emergence of insulin resistance is that of a full train car: if the number of passengers in the train car is low, it is easy to let new passengers in. However, if the train car is full, opening more doors or keeping them open longer will not permit more passengers to enter until some have exited. Therefore, within this framework, the cause of elevated blood glucose is not the inability of insulin-receptor interaction to “open the door”, but the lack of capacity of a cell to store additional glucose.

There exist numerous mathematical models of varying complexity to describe glucose-insulin dynamics ^19^, with the unifying goal of most models to capture emergence of insulin resistance as a function of different internal and external factors. The three best known detailed models of glucose-insulin dynamics are the Hovorka model, the Sorensen model and the UVAPadova model, the latter being accepted by the FDA. These models have very high dimensionality and aim at capturing the many details of glucose-insulin interactions throughout the different tissue compartments in the body in as complete a detail as possible. An excellent summary of the key similarities and differences between these models is given in ^20^. A smaller but still very detailed physiologically-based glucose-insulin model was developed by Alvehag et al. ^21^, where the authors describe feedback control of glucose regulation, keeping track of glucose and the three hormones insulin, glucagon, and incretins, and their distributions through multiple compartments in the body, including brain, liver, heart, lung, pancreas and kidneys. There also have been developed several minimal models of glucose-insulin interactions aimed at answering more focused questions. For example, a recent minimal model with a delay by Murillo et al. ^22^ is used to analyze the impact of plasma free fatty acids on emergence of insulin resistance. One of the keystone minimal models, however, was developed by Bergman et al. ^23^. In this work, the authors were able to quantify the relationships between blood glucose and insulin based on experimental data available at the time; a fascinating account of the development and history of this model can be found in ^24^. A qualitative update to the model has been developed by Topp et al. ^25^, where the authors included emerging understanding of the impact of high blood glucose on insulin-producing Beta cells, which allowed capturing a wider range of physiologically plausible behaviors compared to the original Bergman model; an excellent account of both mathematical and physiological considerations underlying this modification can also be found in ^26^.

In what follows we further update the minimal insulin-glucose model to include the limit on the body’s ability to store excess glucose as a key factor that can affect the emergence of insulin resistance. We show that variation in just one parameter can qualitatively reproduce the Kraft patterns of postprandial glucose-insulin dynamics, capturing the transition between normal glucose-insulin dynamics (Kraft I) to hyperinsulinemia and hyperglycemia, characteristic of Kraft IV (diabetes). We further analyze the relative impact of other factors on change in fasting insulin and glucose levels and show that metabolic environment may determine whether a microscopic tumor might regress or progress. We conclude with a discussion of the emerging evidence that medications developed for managing diabetes could become exciting combination partners to cancer therapy.

### Model description

The proposed model is a modification of the classical models by Bergman et al. ^23^ and Topp et al. ^25^, describing the change over time of blood glucose *G*(*t*), insulin *I*(*t*) and Beta cells B(*t*), with the addition of an equation for intracellular glucose *G*_c_ (*t*). The equation for food *F*(*t*) is added to simulate the effects of an ingested meal on blood glucose and insulin, as one would measure in a standard oral glucose tolerance test (OGTT), to illustrate the ability of the model to qualitatively capture the key patterns of the four Kraft criteria.

It is assumed that a bolus of glucose is absorbed at a rate m, increasing the blood glucose level. Blood glucose can also increase as a result of gluconeogenesis ^27^, where the liver produces and releases glucose during fasting periods. The presence of insulin can additionally inhibit gluconeogenesis ^28^ as a mechanism of tight regulation of homeostatic blood glucose concentrations; these processes are described by the term G_i_(1-pl).

We then assume that glucose can be cleared in an insulin-dependent and insulin-independent manner. Insulin-independent glucose clearance could involve rapid glucose absorption to meet immediate metabolic needs, or it could occur through insulin-independent transporters, such as GLUT1 ^29^. Insulin-dependent glucose clearance involves binding between the insulin molecule and its receptor, which causes the translocation of the GLUT4 transporter on the surface of the cell, allowing the glucose to enter ^30,31^. Here, we describe this process similarly to Bergman et al. ^23^ and Topp et al. ^25^ as *sGI*, where *s* is the rate of insulin-mediated glucose clearance, also often defined as parameter of insulin sensitivity. The ranges for parameter s were chosen based on results reported in Figures 1 and 2 of Kahn et al. ^32^, where the authors report on the inverse relationships between insulin sensitivity parameters and fasting insulin concentrations.

**Figure 1.**
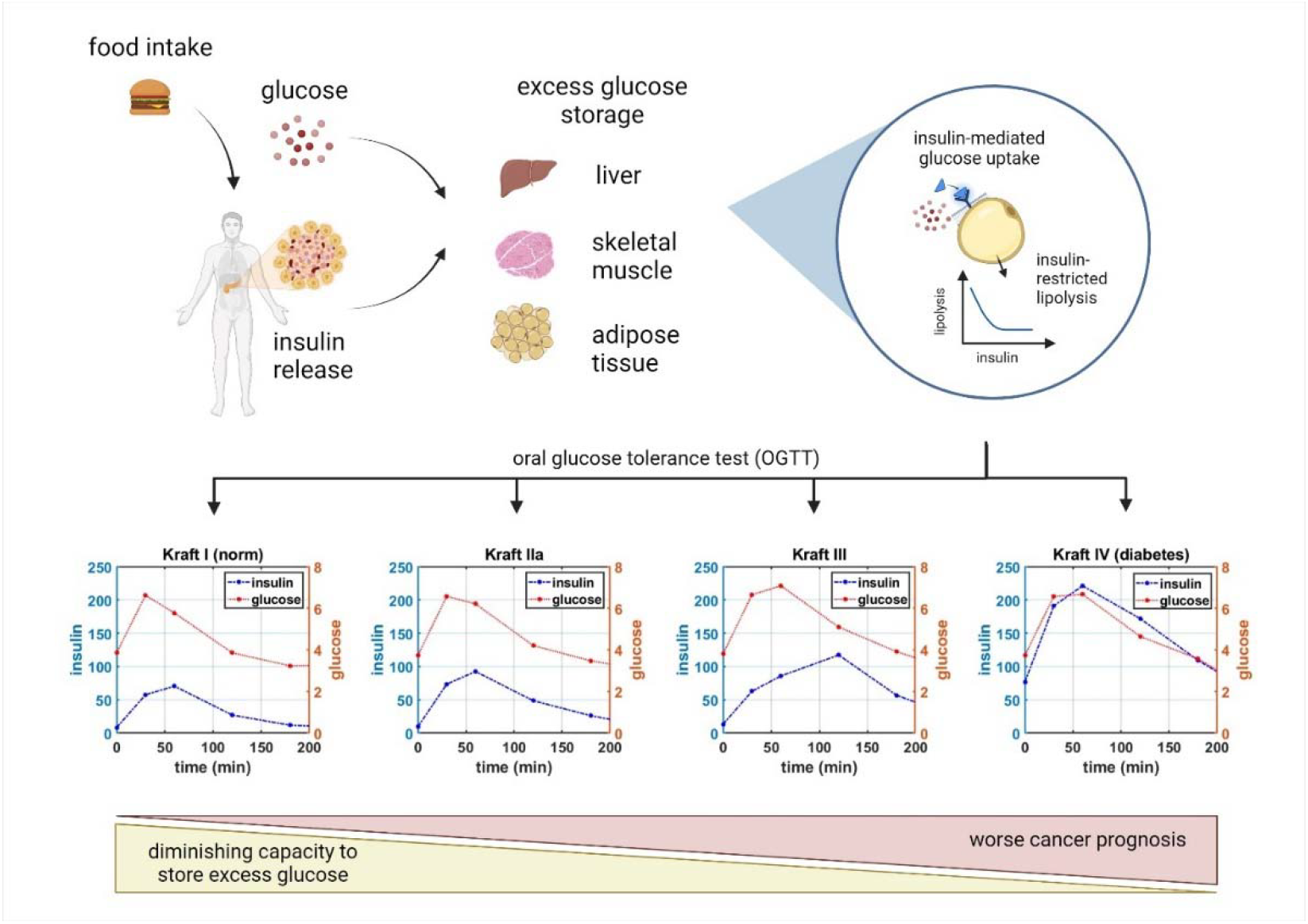
The key processes summarizing glucose-insulin dynamics following glucose ingestion, including insulin-mediated intracellular glucose storage and insulin-restricted lipolysis,which are described in System (1). The model is then used to reproduce four key transitional dynamical regimes, also known as the Kraft patterns ^16^, revealed from a normal response to an oral glucose tolerance test (OGTT) to insulin resistance to diabetic phenotype. The data for the four Kraft criteria were digitized from Crofts et al. ^17^. Glucose is measured in mM; insulin is measured in uIU/L.

**Figure 2.**
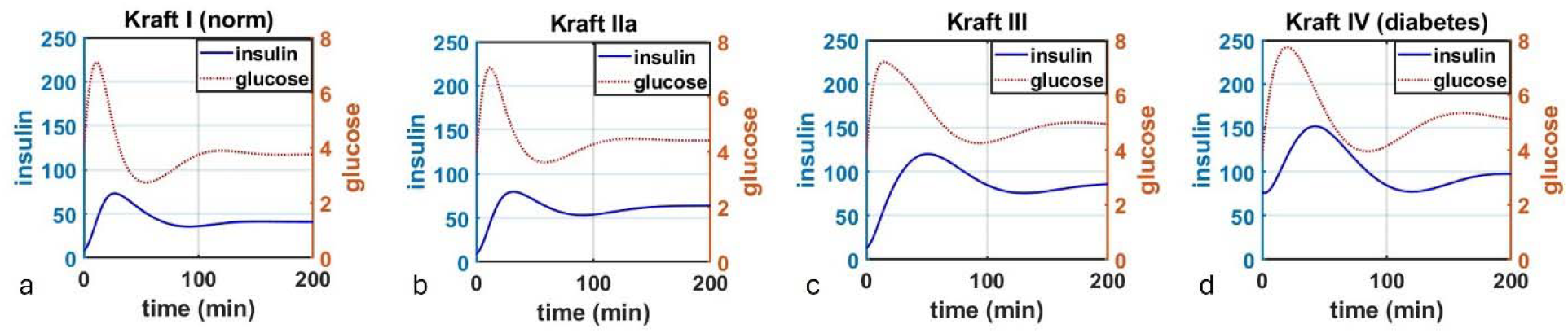
Qualitative simulations of the transition between the Kraft patterns of emerging insulin resistance through variations of parameter K, which represents the capacity to store excess glucose, from (a) *K=50*, to (b) *K=10*, to (c) *K=3*, to (d) *K=0*.*1*.

We then modify this functional form by assuming that extracellular glucose *G*(*t*) is transported into an intracellular glucose storage compartment *G*_*c*_(*t*), which has a carrying capacity k. Excess glucose is assumed to be primarily stored in adipose tissue and skeletal muscle. As a result, the term for insulin-dependent glucose clearance becomes 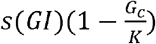.

The equation for intracellular glucose *Gc* then has an insulin-dependent inflow term described above, and insulin-restricted clearance, an assumption that comes from the observations made by Jensen et al. ^33^. In their paper, the authors evaluated the relationship between insulin concentration and lipolysis (fat breakdown). They found that lipolysis was very sensitive to insulin, with higher insulin concentrations resulting in significantly lower lipolysis, while lower concentrations of insulin corresponded to greater fat breakdown. This inverse relationship between lipolysis and insulin is captured by the term 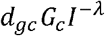, such that larger values of insulin reduce the rate of clearance of intracellular glucose. Parameters d_gc_ and, λ were chosen to qualitatively capture the relationships between lipolysis and insulin in Figure 4 Jensen et al. ^33^ for non-diabetic subjects.

The dynamics of blood insulin and Beta cells are captured similarly to the classic model by Topp et al. ^25^. It is assumed that insulin is produced by Beta cells B(t) in the pancreas. Insulin secretion by is well-described by 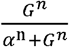, a classical Hill function ^34,35^, which reaches its half-maximal value when glucose concentration is α, and where the steepness of the Hill function is determined by parameter n. In Alcazar and Buchwald ^36^, the authors measured the concentration-response relationship of insulin secretion in both human and murine islets, reporting the slope of the Hill function to be 3.4 for murine and 3.2 for human islets, with half-maximal concentration of glucose being approximately 14 mM for mice vs 8 mM for human cells. As such, parameters n and α in 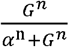 were chosen to be 3.2 and 8, respectively. It is also assumed that insulin is cleared at a normal rate d_l_I.

The equation for the dynamics of Beta cells comes from ^25^, where the authors capture the inverse quadratic relationship between glucose concentration and Beta cell expansion and reduction, a phenomenon known as glucotoxicity. It was quantified experimentally by Efanova et al. ^37^, where rodent beta cell islets were co-incubated with different concentrations of glucose. The authors showed that Beta cell viability increased, up to a point, proportionally to increase in glucose concentrations, presumably to match the necessary insulin production to maintain glucose homeostasis; however, high concentrations of glucose were toxic ^37^. It has been proposed by Karin and Alon ^38^ that such a mechanism exists as a protection against emergence of mutant cells that may perceive low glucose concentrations as higher than they are and compensate by excessive production of insulin, thereby causing potentially fatal hypoglycemia. The simple quadratic equation proposed by Topp et al. ^25^ captures this relationship; parameters a, b, c in Table 1 were taken from Topp et al. ^25^ and converted to corresponding units. Parameter p, describing the Beta-cell dependent insulin secretion rate, was also adapted from Topp et al. ^25^ and converted to corresponding units.

**Table 1.**
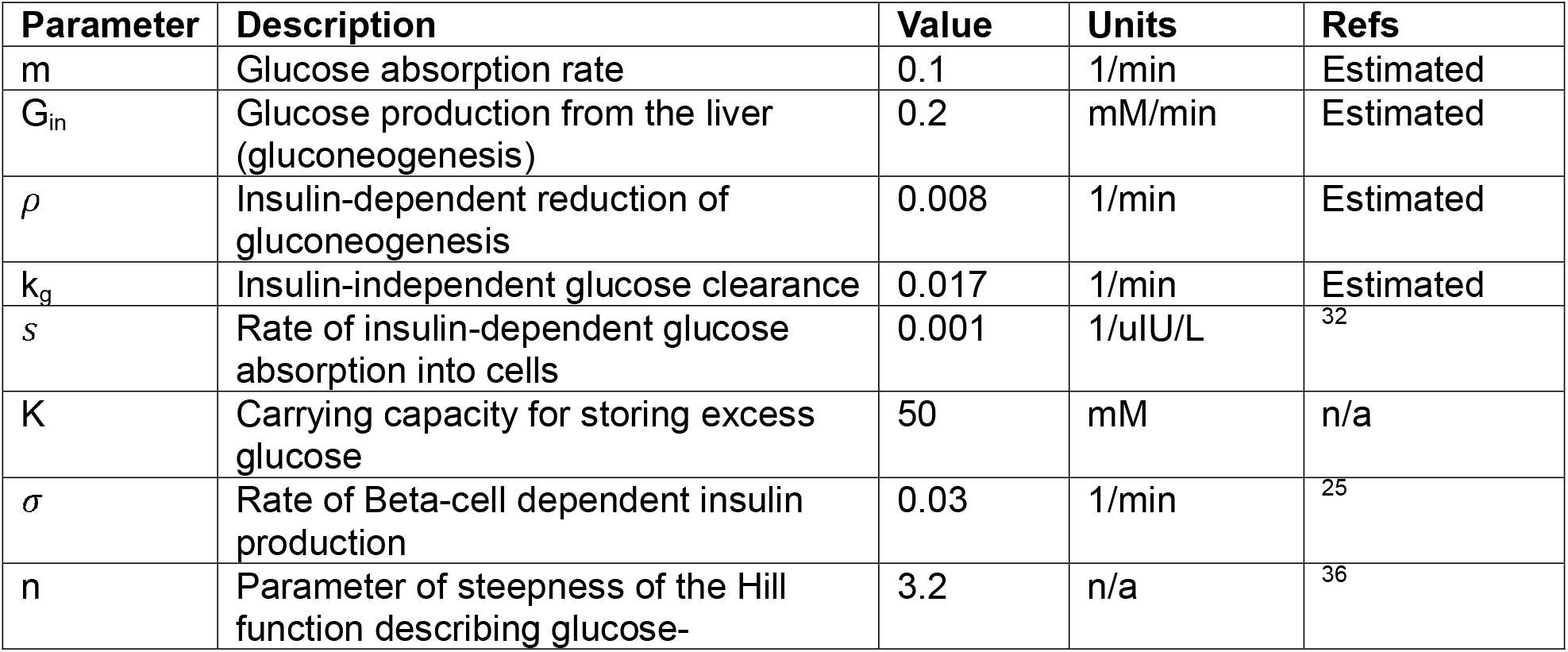
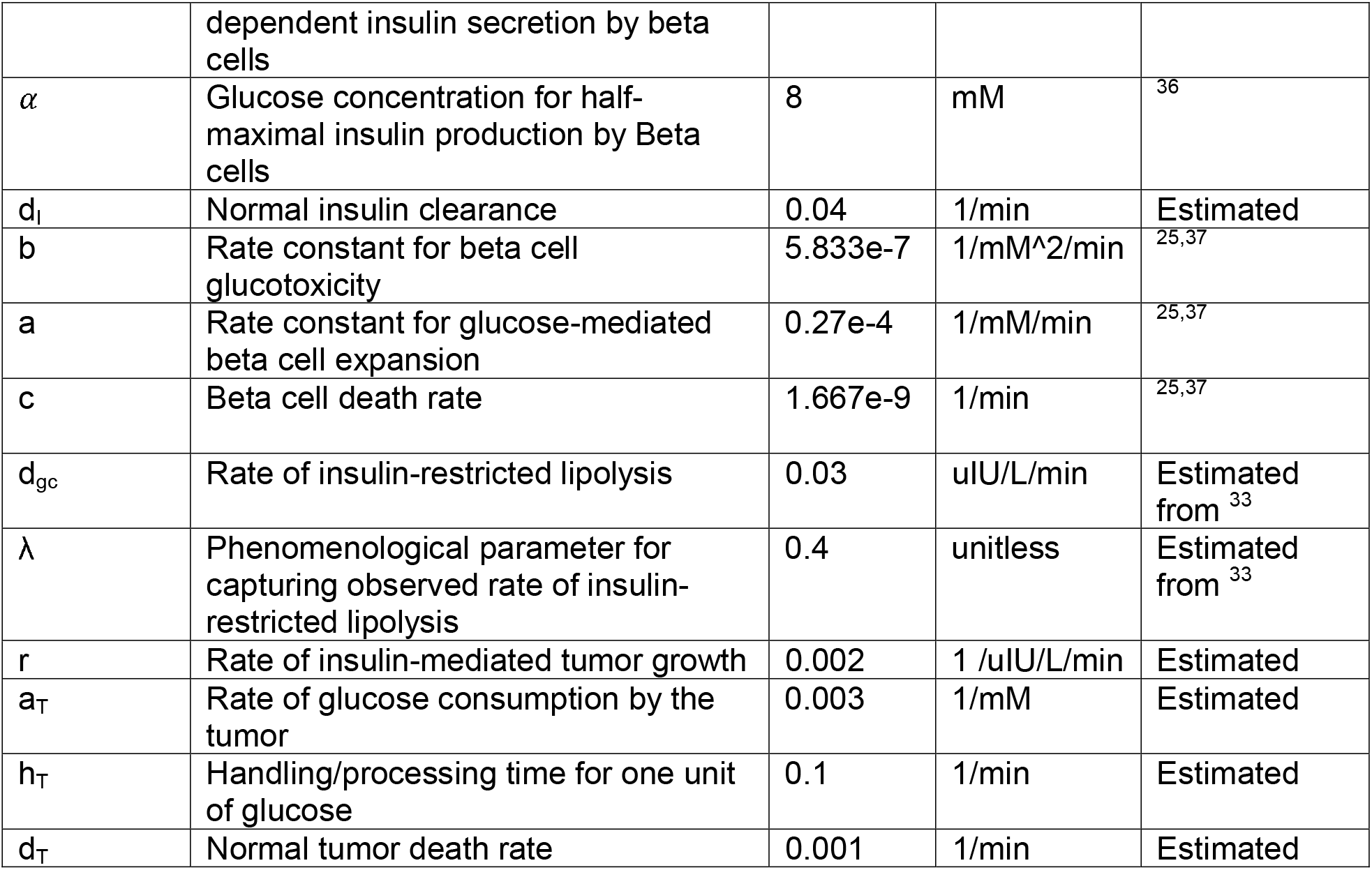
Parameter values used in System (1).

The resulting system of equations is as follows:

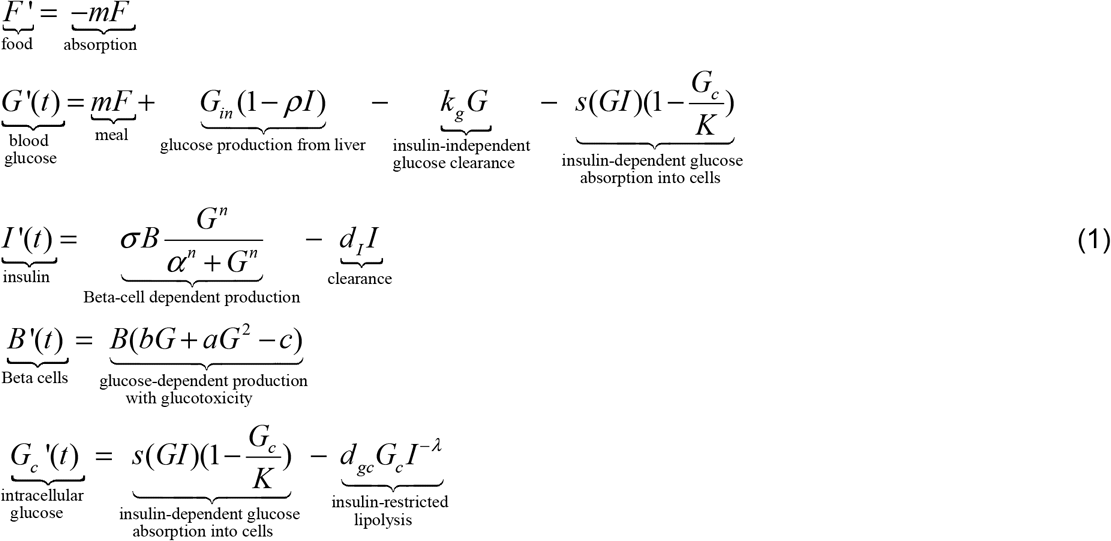

The summary of the key mechanisms described by the proposed model and their connection to the Kraft criteria described in the Introduction is given in Figure 1. Parameter values used in subsequent simulations are summarized in Table 1.

It should be noted that glucose concentrations are typically reported in the literature either in the units of mmol/L, or in the units of mg/dL, with 1 mg/dL of glucose = 18.018 mM. Normal fasting glucose concentrations are typically around 5.5 mM, with normal fasting glucose concentrations of around 100 mg/dL. Insulin data is typically reported either in conventional insulin concentration units (IU/mL) or international units (pmol/L), where 1 uIU/mL = 6.0 pmol/L ^39^. The data reviewed and digitized for this work used both sets of units. Henceforth for consistency, we use mM for glucose and uIU/L for insulin; literature data, when needed, was converted to these units using aforementioned conversion factors.

## Results

The first step of the subsequent analysis is to evaluate whether the model described in System (1) can reproduce the experimentally observed patterns of transition from normal glucose-insulin dynamics (Kraft pattern I) to diabetic (Kraft pattern IV). All subsequent analysis was done in Matlab v2020a using ode23s numerical solver. Parameter values are taken from Table 1 unless indicated otherwise in Figure captions.

### Capturing emergence of insulin resistance through diminishing capacity to store glucose

The classical understanding of the emergence of insulin resistance and subsequent transition to T2D suggests that the key parameter underlying this process is parameter of insulin sensitivity s ^32^; lower insulin sensitivity results in lower value of the term *s*(*GI*), thereby leaving more glucose in the blood. We propose that even for fixed s, it is possible to qualitatively replicate the four main Kraft patterns solely through variation of the parameter of carrying capacity *k* in the updated term 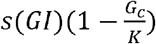 .

In simulations done in Figure 2, we simulated administration of a single bolus of glucose using equation F(t). We varied the value of parameter *k* from *K*=50 (Figure 2a) to *K*=10 (Figure 2b) to *K*=3 (Figure 2c) to *K*=0.1 (Figure 2d). The values of parameter *k* were chosen arbitrarily due to lack of data that would enable its more precise estimate; nevertheless, even with arbitrarily selected value of *k* one can see that diminishing capacity to take up excess glucose from the blood results in transitions from normal insulin-glucose dynamics to both higher baseline levels and longer time needed to return to baseline levels. Furthermore, as one can see particularly in transition from Kraft II (Fig 2b) to Kraft III (Fig 2c), the shape of both curves changes, with a more delayed insulin peak (blue curve) and less rapid decline of glucose (red curve). These results demonstrate that it is possible to capture the features of the Kraft criteria and emergence of insulin resistance solely through progressive decrease of the body’s ability to store excess glucose, particularly in adipose tissue and skeletal muscle, which may in turn arise both from genetic and modifiable factors, as will be discussed in subsequent sections.

Next, we wanted to evaluate the change in equilibrium fasting levels of both glucose and insulin as a function of decreasing carrying capacity *k*. To generate these results, we ran simulations until time t = 2000 (time point chosen arbitrarily to allow the system to fully equilibriate) for different values of *k*, ranging from 100 to 0.1, and collected the corresponding equilibrium values of glucose and insulin. The results are shown in Figure 3. Notably, we highlighted the typical cutoffs for diagnosing pre-diabetes and diabetes for glucose, where normal blood glucose is considered to be <5.5 mM; 5.6-6.9mM is considered to be the pre-diabetes range; over 7.0 mM fasting glucose indicated diabetes ^41^. For insulin, there do not exist such clear cut-off values, with typical values expected to be under 25 uIU/L ^42,43^, although Kraft I criteria suggest normal fasting insulin levels to be <30 uIU/L, with patients meeting Kraft III criteria having fasting insulin up to 50 uIU/L, and Kraft IV criteria patients having fasting insulin >50 uIU/L ^17^.

**Figure 3.**
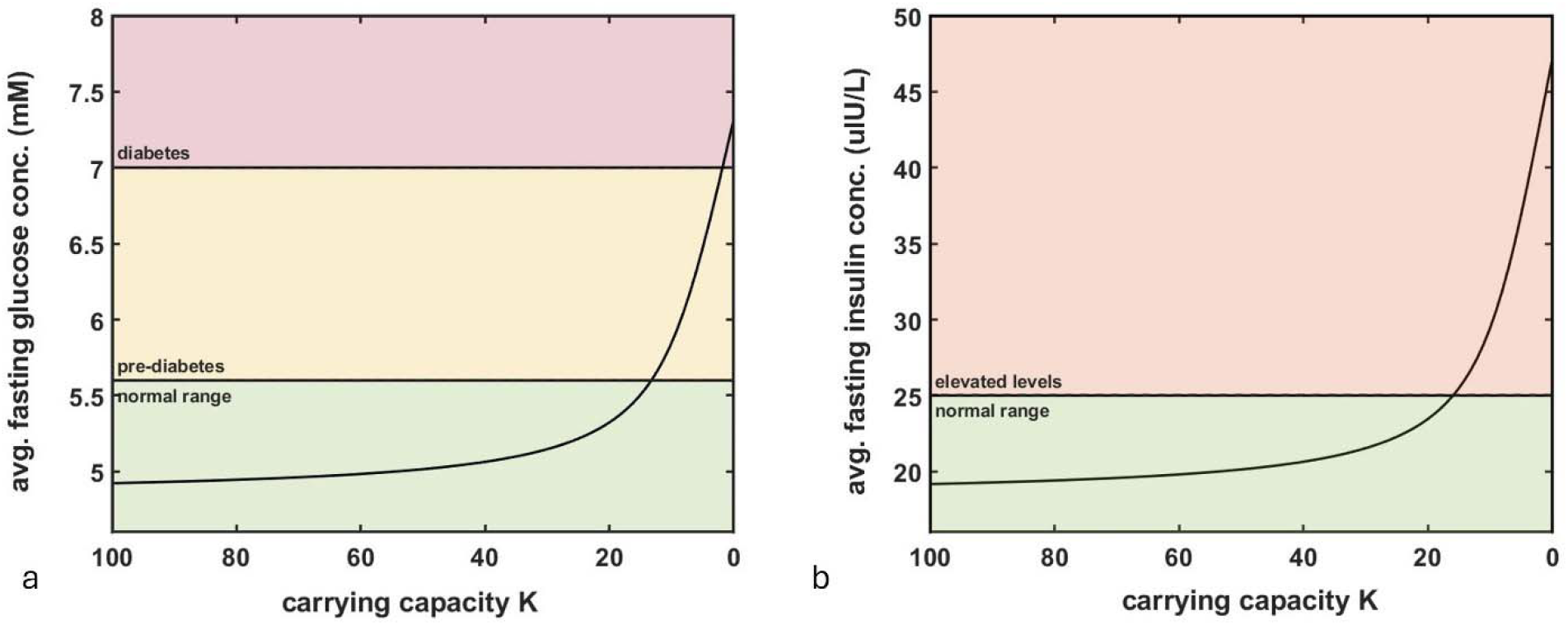
Impact of reduction of the capacity to store excess glucose on fasting levels of (a) glucose and (b) insulin. Reduction in glucose carrying capacity *k* can lead to an increase in average fasting glucose and insulin levels and emergence of a diabetic phenotype.

As can be seen in Figure 3, assuming a maximum value of *K = 100*, decreasing its value starts to impact baseline fasting levels of glucose and insulin only around when only 15-20% of capacity to store excess glucose remains, which indicates that there exists a large buffer allowing the body to maintain normal levels of glucose and insulin even as *k* decreases.

Next, we wanted to evaluate the impact of variation of both parameter *k* and the parameter of insulin sensitivity *s* on the average fasting values of both insulin and blood glucose. Parameter *k* was varied from 0.01 to 40, since analysis reported in Figure 3 revealed that larger values of *k* are not expected to affect glucose and insulin fasting levels; parameter *s* was varied from 0.0001 to 0.002 to fall within the range of values for insulin sensitivity reported in Kahn et al. ^32^. Similar to the previous set of simulations, we varied parameters *s* and *K* and reported the corresponding fasting levels of glucose and insulin. As one can see in Figure 4, at very low values of *s* (extrermely low insulin sensitivity), even high capacity to store excess glucose does not protect from elevated fasting glucose and insulin. However, for larger values of *s* (normal insulin sensitivity), parameter *K* plays a key role. Together, these results add another dimension to understanding the impact of insulin sensitivity on fasting glucose and insulin levels: even when one’s insulin sensitivity is high, it is still possible to develop insulin resistance, hyperglycemia and hyperinsulenemia through diminishing ability to store excess glucose.

**Figure 4.**
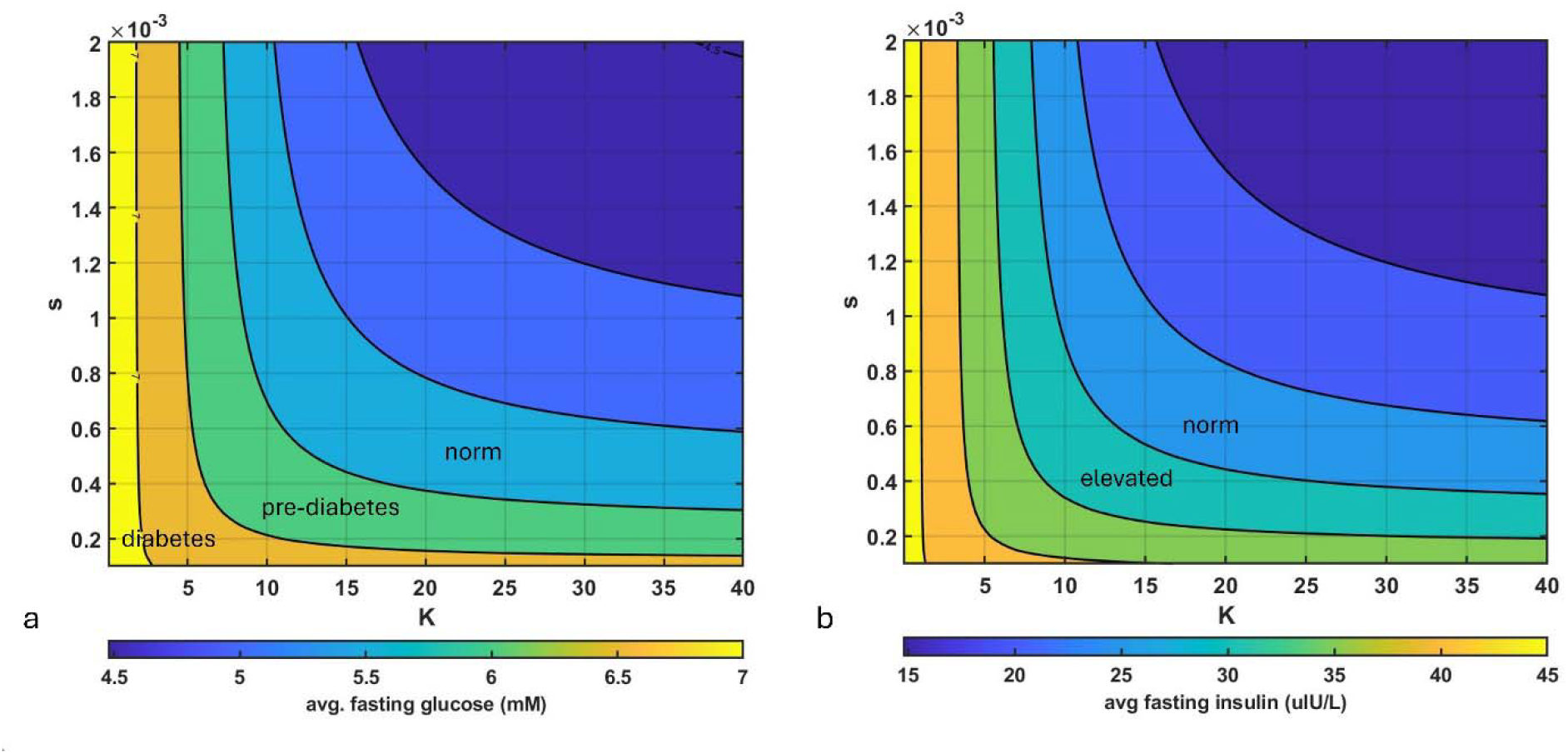
Impact of variation of change in both the capacity to store excess glucose (parameter K) and insulin sensitivity (parameter s) on fasting (a) glucose and (b) insulin levels. High capacity for storing excess glucose can to an extent compensate for low insulin sensitivity; reduction of the capacity to store glucose can lead to accelerated development of a diabetic phenotype as measured by fasting levels of glucose and insulin.

### Impact of hyperinsulinemia on tumor growth

In Hopkins et al. ^10^, the authors demonstrated that hyperinsulinemia was the driver of emergence of resistance to PI3K inhibitors in mouse cancer models regardless of etiology; furthermore, they showed that reversing hyperinsulinemia was sufficient to restore sensitivity to the drug for all twelve tested cancer cell lines. They proposed that the underlying mechanism for their observations was not elevated glucose acting as additional fuel but instead elevated insulin acting as an accelerant of tumor growth.

To assess the impact of hidden hyperinsulinemia on tumor growth, we add an additional equation for a tumor that grows both proportionally to available glucose and insulin:

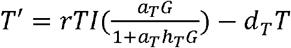

Here, r is the tumor growth rate, *a*_*T*_ is glucose consumption rate, *h*_*T*_ is glucose handling rate (i.e., the amount of time it requires to metabolize it) ^44^, and *d*_*T*_ is intrinsic tumor death rate. We next simulated tumor growth over time for an individual with glucose and insulin levels corresponding to Kraft I, II, III and IV as shown in Figure 2.

As one can see in Figure 5, the same tumor can grow or regress depending on the metabolic state of its host. In this set of simulations, the tumor will regress for an individual with Kraft I or II metabolic profile, since the death term 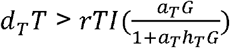. However, for Kraft III and IV, due to higher levels of insulin and glucose, cumulative growth term 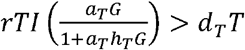, resulting in net tumor growth.

**Figure 5.**
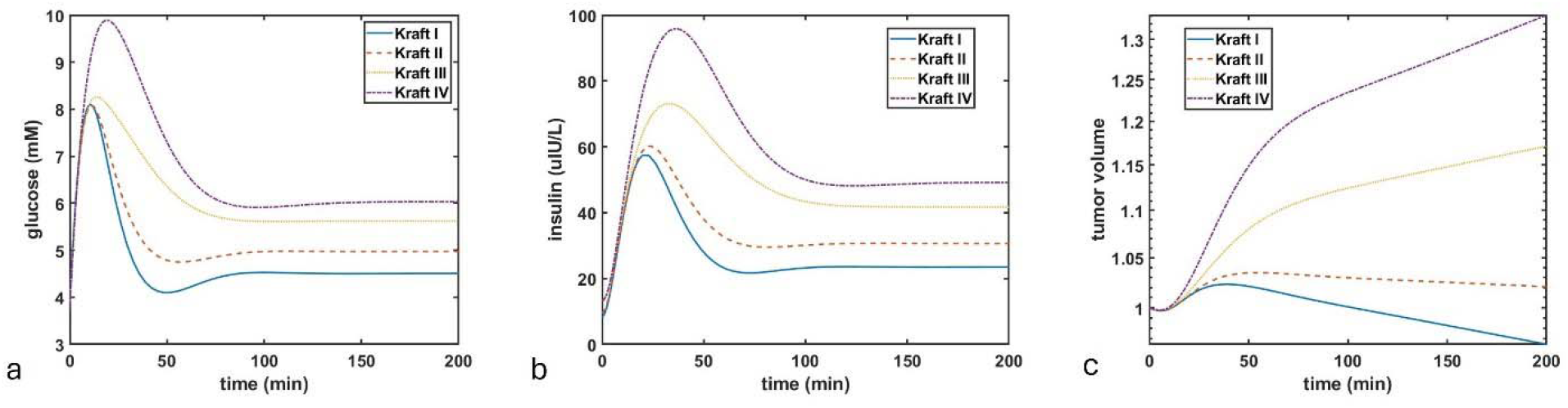
Projected impact of the metabolic environment of the host as defined by Kraft patterns following an OGTT on subsequent dynamics of a microscopic tumor. Kraft patterns are modeled through variation of parameter K, with *K = 50* for Kraft I, *K = 10* for Kraft 2, *K = 3* for Kraft 3, and *K=0*.*1* for Kraft IV, with corresponding changes over time for (a) glucose, (b) insulin, and (c) volume of a microscopic tumor.

Next, we simulated the impact of parameters *s* and *K* on whether a microscopic tumor might progress or regress. For this, we once again varied parameter *K* from 0.01 to 40, and parameter *s* from 0.0001 to 0.002. We ran the simulation till *t = 2000* and calculated whether the difference between the initial tumor size and its size at *t = 2000* was positive (tumor progression) or negative (tumor regression). As one can see in Figure 6, there exists a region of low insulin sensitivity (which, as was seen in Figure 4 above, corresponds to regions of higher fasting glucose and insulin levels), where, should a tumor appear, it is expected to progress. However, for individuals with higher insulin sensitivity, larger capacity to store excess glucose and therefore lower fasting glucose and insulin levels can prevent an emerging tumor from progressing.

**Figure 6.**
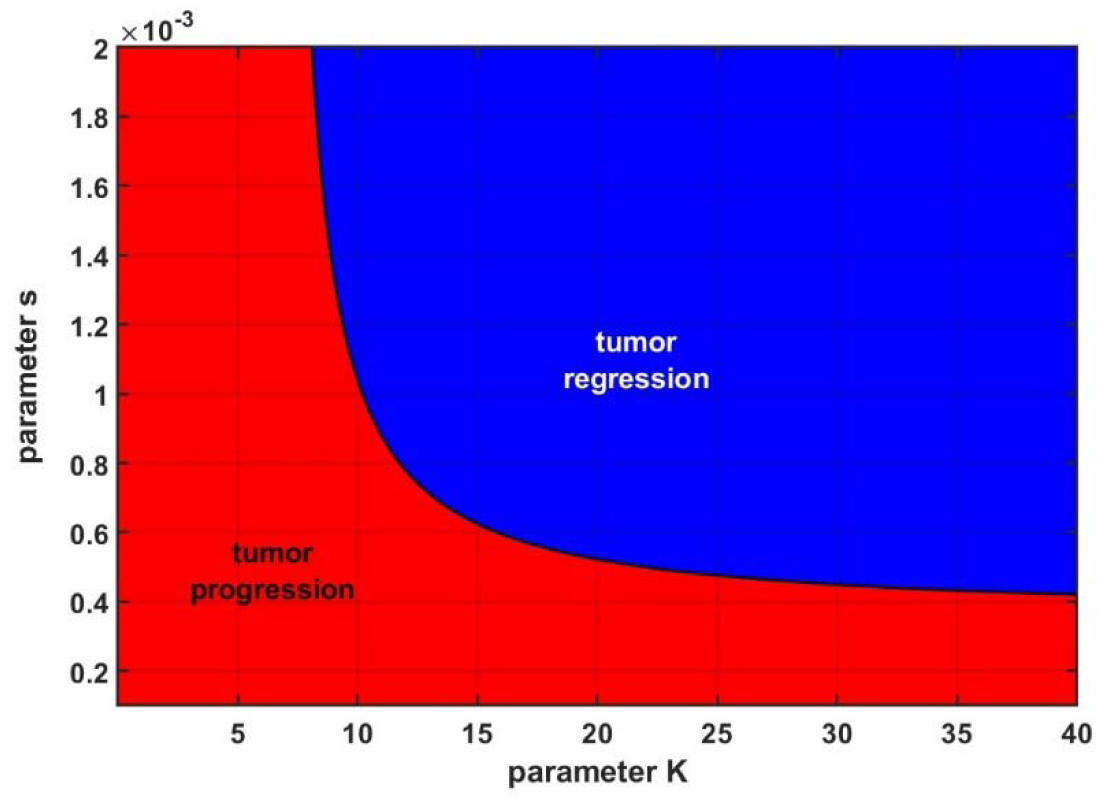
Projected impact of fasting the metabolic environment of whether a tumor would progress (red) or regress (blue) as a function of both the capacity to store excess glucose (parameter K) and insulin sensitivity (parameter s).

While the parameters for tumor growth and death rates in this model were chosen arbitrarily, the simulations nevertheless serve to illustrate how hidden hyperinsulinemia can act to promote the growth of a microscopic tumor that would not thrive in a more metabolically healthy environment.

## Discussion

Emergence of insulin resistance and subsequent hyperinsulinemia frequently occur in individuals with normal fasting glucose ^16,17^; however, chronic hyperinsulinemia that accompanies hyperglycemia, characteristic of T2D, can contribute to the elevated risk of cardiovascular disease, dementia and cancer ^7,8,45,46^. Classical understanding of emergence of insulin resistance assumes loss of functional signaling between insulin and its receptor and thus a lower sensitivity to insulin signaling ^32^. Here we describe an updated framework of describing glucose-insulin dynamics, where emergence of insulin resistance that precedes hyperglycemia characteristic of T2D is a result not of loss of sensitivity to insulin but of diminishing capacity to store excess glucose (Figure 1).

The implications of this framework are studied using a modification introduced into the well-established minimal models of glucose-insulin dynamics, developed by Bergman et al. ^23^ and Topp et al.^25^. We demonstrate that reduction of the carrying capacity parameter *K* is sufficient to qualitatively describe both the key features of the four key patterns ^16,17^ of glucose-insulin dynamics during a standard OGTT (Figure 2), and the incremental increases in fasting levels of both glucose and insulin, capturing the transition from norm to pre-diabetes to the T2D range with a single parameter (Figure 3). We demonstrate that even for individuals with high insulin sensitivity, it is still possible to develop insulin resistance, hyperglycemia and hyperinsulinemia through diminishing ability to store excess glucose.

Finally, we build upon the observations of Hopkins et al. ^10^, where the authors showed that development of diabetic phenotype in mouse cancer models resulted in emergence of reversible therapeutic resistance to PI3K inhibitors due to growth signals from elevated insulin overwhelming the cytotoxic effects of the drug. Using the proposed model, we demonstrate that change in metabolic environment and increased fasting glucose and insulin can enable the microscopic tumor to transition from regression to progression (Figures 5). This prediction is consistent with observations that pre-existing T2D is associated with worse prognosis for many cancer types ^9,47,48^. Furthermore, our results suggest that hidden hyperinsulinemia, which can occur independently of T2D and in individuals with normal glucose tolerance, which was the case in over half (n=4185) of participants in the analysis done by Crofts et al.^17^, can prime the environment for tumor progression, should a cancer arise. Finally, we demonstrated that change in one’s ability to store excess glucose can be an important modifiable factor in promoting cancer regression vs progression (Figure 6).

Excess glucose can be stored in three main compartments: adipose tissue, liver, and skeletal muscle. The capacity for how much excess glucose can be stored in the adipose tissue is likely in part genetic, with some individuals having larger carrying capacity, allowing them to accumulate fat tissue without acquiring a diabetic phenotype, while others may have a smaller carrying capacity, and thus begin to exhibit elevated glucose and insulin levels even in the absence of high body fat ^49,50^. Interestingly, in an elegant experiment, Gavrilova et al. ^51^ showed that surgical transplantation of wild type adipose tissue into severely diabetic lipoathropic mice reversed the diabetic phenotype by effectively increasing the adipose carrying capacity and creating a compartment, into which excess glucose could be stored.

Storing too much excess glucose as fat in the liver can lead to the development of non-alcoholic fatty liver disease (NAFLD), which currently has a global prevalence of approximately 25% ^52^. In the early stages, NAFLD is fully reversible ^53,54^. However, if it is left unchecked, it can lead to non-alcoholic steatohepatitis (NASH), which in turn can lead to cirrhosis; at the current rate, cirrhosis with NASH is poised to become the leading indication for liver transplantation in the next decade ^55,56^.

Finally, the largest storage depot for excess glucose is skeletal muscle, which is also the most modifiable factor for varying carrying capacity *K*. It can be increased through exercise, or lost through sedentary lifestyle ^57,58^, as a result of the normal aging process ^59^, or even as a result of bed rest, particularly for older adults ^60^. Interestingly, exercise additionally enables activation of insulin-independent glucose clearance mechanisms, a strategy that has been used by T1D patients to reduce the need for exogenous insulin ^61,62^.

Advances in glucose monitoring technology have enabled understanding of the dynamics of glucose not only during an OGTT but throughout the day, which has been an extremely important tool for blood sugar management for both T1D and T2D patients. Continuous glucose monitors (CGMs) are small devices with a thin membrane-coated wire that inserts into subcutaneous tissue. After coming in contact with interstitial fluid, they generate electrochemical signals, which are received by a transmitter on top of the sensor, where an algorithm converts the electrochemical signal to a glucose value ^63^. Notably, the ability to measure glucose in the interstitial space using an enzymatic method, which does not require multiple washes, allows for the continuous measurement of glucose in these devices. Unfortunately, measuring insulin requires using either ELISA or radioimmune assays ^64^, thereby currently making immediate readings impossible. Nevertheless, even in the absence of continuous insulin monitors, it may be possible to monitor postprandial glucose levels within the frameworks of Kraft patterns.

Pharmacologically, until recently, typical treatments for T2D involved medications aimed at increasing insulin production from pancreas, such as sulfonylureas ^65^, as well as administration of exogenous insulin when the capacity of the pancreas to produce insulin is pharmacologically maximized. In the last two decades, a new class of drugs for normalizing blood glucose has emerged. Of particular note are sodium glucose cotransporter 2 (SGLT2) inhibitors, which block the activity of co-transporter protein responsible for up to 90% of glucose reabsorption in kidneys. As a result, blockade of SGLT2 increases renal excretion of glucose without additional insulin, thereby systematically lowering the amount of excess glucose in the blood without the need to store it in the body ^66,67^. Furthermore, SGLT2 is expressed in certain types of cancer ^68,69^, making it a potential target for cancer therapy as well. Indeed, in their observational study that included nearly 25000 patients diagnosed with non-small cell lung cancer (NSCLC) between 2014-2017 who had a pre-existing diagnosis of T2D, Luo et al. ^70^ assessed whether taking SGLT2 affects NSCLC prognosis and survival. Among patients who had used an SGLT2 inhibitor for over 12 months, the authors showed a 46% relative reduction in all-cause mortality compared to non-users. Whether the benefit came solely from blocking the mechanism of action of SGLT2, affecting cancer cell metabolism, or from the systemic lowering of glucose and insulin, remains to be evaluated. If so, then some of the drugs that act to normalize both blood glucose and insulin might be exciting future combination partners for cancer therapy.

## Data Availability

N/A

